# New method for high-throughput measurements of viscosity in submicrometer-sized membrane systems

**DOI:** 10.1101/736785

**Authors:** Grzegorz Chwastek, Eugene P. Petrov, James Peter Sáenz

## Abstract

In order to unravel the underlying principles of membrane adaptation in small systems like bacterial cells, robust approaches to characterize membrane fluidity are needed. Currently available relevant methods require advanced instrumentation and are not suitable for high throughput settings needed to elucidate the biochemical pathways involved in adaptation. We developed a fast, robust, and financially accessible quantitative method to measure microviscosity of lipid membranes in bulk suspension using a commercially available plate reader. Our approach, which is suitable for high-throughput screening, is based on the simultaneous measurements of absorbance and fluorescence emission of a viscosity-sensitive fluorescent dye DCVJ incorporated into a lipid membrane. We validated our method using artificial membranes with various lipid compositions over a range of temperatures and observed values that were in good agreement with previously published results. Using our approach, we were able to detect a lipid phase transition in the ruminant pathogen *Mycoplasma mycoides*.

---

Viscosity is a crucial physical property of living membranes that is tightly regulated though homeostatic adaptation to environmental and physiological challenges.^[1, 2, 3, 4, 5, 6,7]^ Measuring viscosity is important for investigating the mechanisms involved in membrane adaptation and to constrain the range of membrane properties that can support life. In particular, studying relatively simple bacterial model organisms can provide insight into fundamental principles underlying membrane homeostasis and adaptivity. Presently, however, there are no high throughput or broadly accessible methods to measure viscosity in bacterial cells or submicron scale synthetic membrane systems.

Currently existing methods for measuring membrane viscosity are relatively low throughput or require specialized instrumentation not available to many laboratories. Fluorescence correlation spectroscopy (FCS) can provide estimates of diffusivity of a molecular probe.^[8,9]^ FCS, however, requires a relatively specialized microscopy setup, and measuring diffusion in submicrometer scale membrane systems can be particularly challenging. Similarly, fluorescence recovery after photobleaching is not feasible on small vesicles because of spatial resolution limitations.^[10]^ Another common method to estimate membrane viscosity is based on emission anisotropy of fluorescent probes.^[11]^ In this case, however, the interpretation of results is not always straightforward.^[12]^ Recently, there have been a number of promising studies measuring viscosity utilizing the fluorescence lifetime of viscosity-sensitive fluorescent probes. While this approach could be applied to submicron membrane systems, the technology is fairly expensive, and high throughput instrumentation is currently not commercially available. In this study, we developed a method to estimate membrane viscosity by measuring the relative brightness of a viscosity-sensitive fluorescence dye using a simple plate reader capable of simultaneously measuring absorbance and fluorescence emission. Our method is based on the empirical finding of Förster and Hoffmann that certain fluorescence probes undergoing twisted intramolecular charge transfer (TICT) show a power-law dependence of their brightness (fluorescence quantum yield, *ϕ*) on the viscosity *η* of bulk solvents *ϕ* ∝ *η*^*p*^.^[13]^ This relation holds in cases where the non-radiative decay rate is controlled by the viscosity of the medium, as long as the non-radiative decay rate is much higher than rate of radiative decay rate. In that case one can use this effect to monitor the microviscosity via either fluorescence quantum yield or by the excited-state lifetime.^[14, 15, 16, 17, 18]^

Among others, 9-(2,2-dicyanovinyl)julolidine (DCVJ) is a well characterized TICT probe (Fig. 1) with the fluorescence emission in the visible range of the light spectrum.^[19]^ It was shown that DCVJ can be used to estimate the viscosity in lipid membranes using the relative quantum yield approach.^[16]^ Moreover, the dye is commercially available at a very accessible price making it a perfect candidate for application in large-scale screening assays.

**Figure 1.**
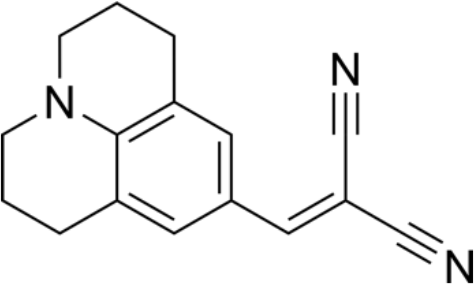
Chemical structure of 9-(2,2-dicyanovinyl)julolidine (DCVJ).

In order to estimate membrane viscosity, we compared the relative brightness of DCVJ incorporated into liposomes with the brightness of DCVJ measured in solvents of known viscosities. To this end, we developed an experimental protocol that overcomes the limitations of the analytical noise of the plate reader and artifacts related to the sample structure and construction of the multi-well plate. To validate our method, we measured the relative brightness of DCVJ in liposomes composed of several well-characterized lipid species and lipid mixtures at the physiological range of temperatures. We showed that the viscosity activation energies obtained using our method agree within experimental error with those reported in the literature. Using our approach, we report values for membrane viscosity in liposomes made of several different lipid species and provide a first estimate of the membrane viscosity of the ruminant pathogen *Mycoplasma mycoides*. The method we report provides an affordable and fast means to measure the viscosity of membranes and could be used in screening settings where, e.g., a large number of bacterial strains or mutants could be studied.

## Results and Discussion

### 1. Establishing a plate reader assay for measuring variations in membrane viscosity

We aimed at establishing a method that would allow for accurate measurements of the membrane viscosity in lipid vesicles. To this end, we made use of the power-law dependence of the fluorescence quantum yield of DCVJ on the viscosity of its microenvironment. As the measurement of the quantum yield is problematic in our experimental setting, we replace it by the fluorescence brightness denoted here as *R* and defined as the ratio of integrated fluorescence DCVJ emission and the absorbance of the probe in the sample (for details, see Experimental Section). To calibrate our method, we carried out measurements of *R* DCVJ in several media covering a wide range of viscosities expected for lipid membranes. To this end, we used two neat viscous solvents, glycerol and ethylene glycol, whose viscosity strongly depends on the temperature, and measured *R* in these solvents over a range of temperatures relevant for our membrane experiments. Additionally, *R* was measured in a series of glycerol-methanol mixtures at the room temperature. The results of the temperature-independent and temperature-dependent measurements agree with each other very well (Fig. 2), which shows that the fluorescence brightness of DCVJ can be used to report the viscosity of its microenvironment, irrespective of the temperature. As expected, *R* shows a power-law dependence on the viscosity with the exponent *p* = 0.53 ±0.01 falling into range of values (0.51-0.59) reported previously.^[15]^ This power-law correspondence allows us to convert the measured brightness of the DCVJ fluorescence into the viscosity of its microenvironment (for details, see Experimental Section). It is important to emphasize that the fluorescence quantum yield — and hence fluorescence brightness *R* — of a molecular rotor reflects the rotational mobility of the dicyanovinyl moiety of the molecule. As a result, the method reports the viscosity of the membrane microenvironment of the probe, which will be referred in what follows as microviscosity. In contrast, the methods based on translational diffusion of relatively large membrane inclusions like proteins, colloidal particles, or membrane domains, give information on the surface viscosity of the lipid bilayer,^[20, 21, 22]^ which, with the use of the bilayer thickness, can be converted into an estimate of the bulk viscosity of the membrane material. In contrast, methods based on measuring translational diffusion of fluorescent lipid analogs or fluorescently labeled lipids, which are too small to warrant the hydrodynamics-based description of their motion, do not allow one to obtain valid estimates of membrane viscosity.

**Figure 2.**
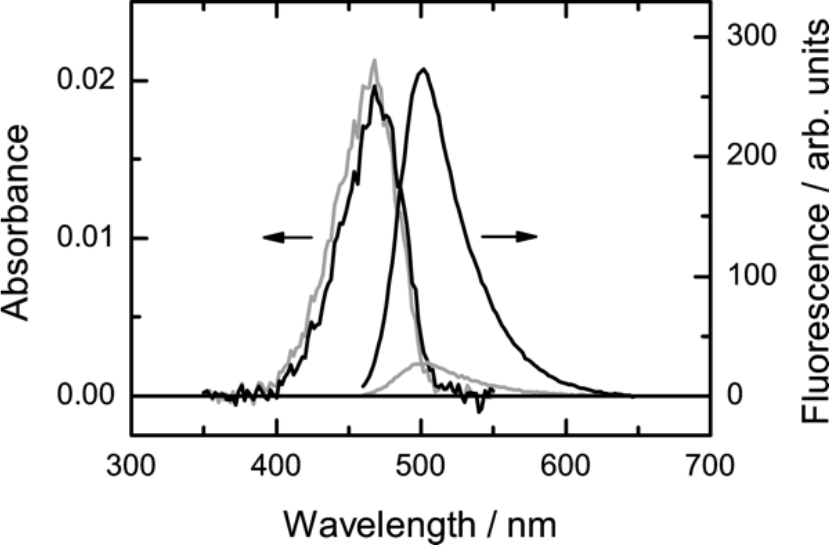
Absorbance and fluorescence emission spectra of DCVJ in glycerol and ethylene glycol at 18 °C. At the same dye concentration, absorbance spectra are virtually identical. In contrast, fluorescence emission depends strongly on the viscosity of the solvent, and under the same experimental conditions, a substantial increase in the fluorescence intensity is seen in more viscous glycerol.

First, we tested the fluorescence response of DCVJ to temperature in membranes comprised of DOPC which is an unsaturated phospholipid with a melting temperature below −20 °C. Thus, under our experimental conditions it is in the fluid state far from the lipid melting phase transition. The DOPC viscosity estimated from the DCVJ fluorescence brightness for the temperature range used in our experiment could be very well described by the Arrhenius law (Fig. 3) with the activation energy of 54 ± 9 kJ/mol, which agrees with previous findings based on measurements of the fluorescence lifetime of a molecular rotor (Table 1). The previously reported absolute values of the viscosity are in the range of 13 to 74 mPa·s and are of the same order of magnitude as reported by Kung and Reed for the DPPC membrane in a liquid phase.^[23]^ While it is clear that methods reporting microviscosity are relative, it is still valuable to estimate the scale of discrepancy between the relevant methods. When compared to membrane studies involving the lateral diffusion of membrane inclusions fulfilling the requirements of the hydrodynamic model for membrane diffusion,^[20, 21, 22]^ our results are roughly a factor of 3 lower (128 mPa·s at 24 °C vs 41 ± 10 mPa·s at 25 °C in our case). On the other hand, Wu et al. using an approach based on the microviscosity dependence of the fluorescence lifetime of a molecular rotor reported 228 mPa·s for DOPC membrane at 25 °C. Hence, it is clear that discrepancies of a factor of 2 to 6 are common for such measurements and show the specificity of the approach rather than its drawbacks.

**Table 1.** Arrhenius activation energies characterizing the viscosity and Brownian motion in lipid membranes of various compositions.

| Membrane composition | $E_a$, kJ/mol | Method | Lipid system | Quantity | Reference |
| --- | --- | --- | --- | --- | --- |
| DOPC | 54 ± 9 | fluorescence brightness | LUVs | viscosity | this work |
|  | 27 | pfg NMR | supported lipid multibilayers | diffusion coefficient | 25 |
|  | 46.1 | fluorescence lifetime | LUVs | viscosity | 18 |
| POPC | 53 ± 10 | fluorescence brightness | LUVs | viscosity | this work |
|  | 28 | pfg NMR | supported lipid multibilayers | diffusion coefficient | 25 |
|  | 48.6 | fluorescence lifetime | LUVs | viscosity coefficient | 18 |
| DOPC/chol 6:4 | 63 ± 8 | fluorescence brightness | LUVs | viscosity | this work |
|  | 32 | pfg NMR | supported lipid multibilayers | diffusion coefficient | 25 |
| SOPC | 68 ± 8 | fluorescence brightness | LUVs | viscosity | this work |
|  | 31 | pfg NMR | supported lipid multibilayers | diffusion coefficient | 25 |
| DLPC | 52 ± 13 | fluorescence brightness | LUVs | viscosity | this work |
| DPPC/chol 1:1 | 54 ± 11 | fluorescence brightness | LUVs | viscosity | this work |
| POPC/chol 1:1 | 58 ± 12 | fluorescence brightness | LUVs | viscosity | this work |
|  | 37 | pfg NMR | supported lipid multibilayers | diffusion coefficient | 25 |

**Figure 3.**
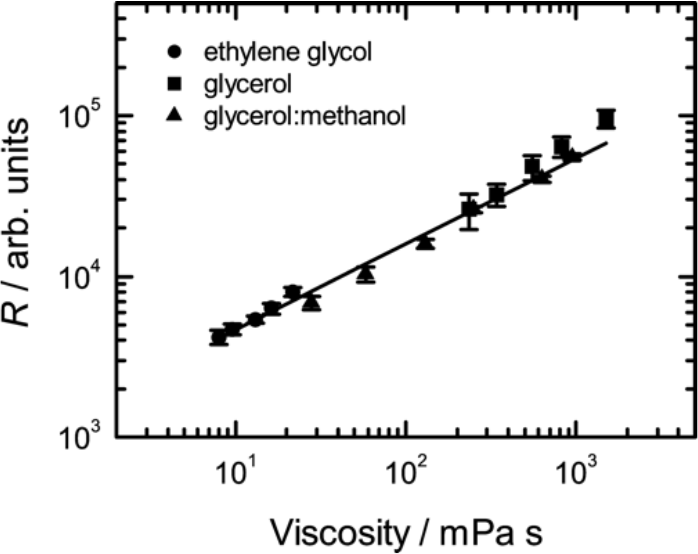
Relative brightness of DCVJ in solvents of different viscosity. Viscosity of glycerol and ethylene glycol was varied by changing the temperature. Mixtures of glycerol and methanol at various ratios were measured at constant temperature (22 °C). The line shows the power law fit.

### 2. Influence of cholesterol on viscosity of phospholipid membranes

Living organisms use cholesterol to control and adapt their membranes to constantly changing environmental conditions. Therefore, a robust assay to estimate the influence of cholesterol on membrane properties is crucial. To test the sensitivity of our method to cholesterol content, we measured the viscosity of DOPC membrane supplemented with 40 mol% of cholesterol. (Fig. 4). Cholesterol introduced a substantial increase in the viscosity at all temperatures (Fig. 4). The estimated activation energy for viscosity is 63 ± 8 kJ/mol, roughly 17 % higher than that for the pure DOPC membrane. Our value is thus close to that of Petrov and Schwille who analyzed results of Cicuta et al.^[22, 24]^ for the liquid disordered (Ld) phase of DOPC/DPPC/cholesterol ternary mixture, for which the activation energy was estimated to be 77 kJ/mol. A similar increase in the activation energy (18 %) upon addition of the same amount of cholesterol was found by Filippov et al. using an NMR-based approach;^[25]^ here, however, it is important to point out that the activation energies were reported not for membrane viscosity, but rather for the diffusion coefficient of a deuterated lipid in the membrane, which could potentially explain the difference in the results. Based on fluorescence lifetime measurements of a molecular rotor, Wu et al. have studied the lipid mixture, and, while the activation energy of the viscosity was not reported,^[18]^ they found a relatively moderate increase (16 %) in the absolute viscosity. In contrast, we observed an 85 % increase in the membrane viscosity upon addition of cholesterol for the same temperature. A similar trend in viscosity was observed in experiments involving translational diffusion of membrane inclusions that fulfil the requirements of the hydrodynamic model: based on the published results on the surface membrane viscosity obtained there, we calculated the bulk membrane viscosity for DOPC (140 mPa·s, 24 °C) and cholesterol-enriched Ld phase of DOPC (270 mPa·s, 25 °C) assuming the membrane thickness to be 3.7 nm.^[26, 22, 27]^ The viscosity difference between pure lipid and cholesterol-doped bilayer comprises 70 % and is reasonably close to our results. Therefore, we argue that the sensitivity of our approach to cholesterol is similar to that of studies based on translational diffusion of membrane inclusions and thus correctly reflects the viscous properties of the membrane material.

**Figure 4.**
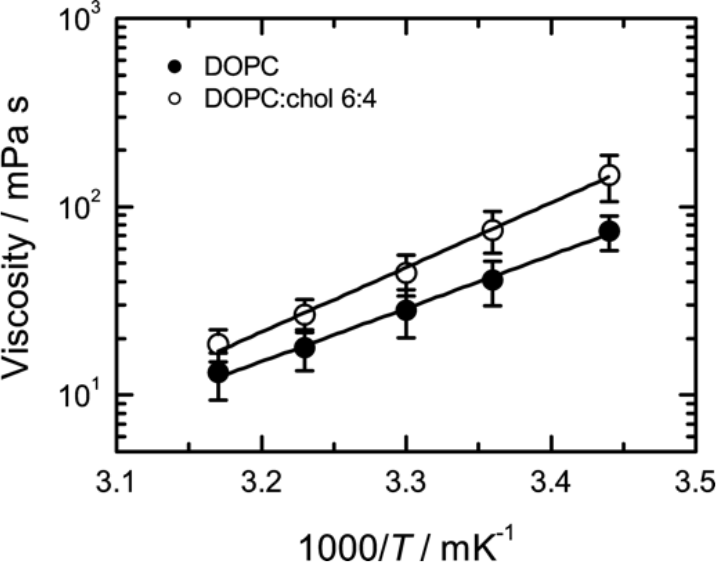
Viscosity of lipid membrane composed of DOPC and DOPC/cholesterol 6:4 (mol:mol). Viscosity data are shown with a fit using the Arrhenius law (line). The activation energies were 54 ± 9 and 63 ± 8 kJ/mol for pure and cholesterol doped DOPC membrane, respectively.

### 3. Sensitivity to variation in phospholipid acyl chain saturation and length

Besides cholesterol, the physical properties of biological membranes can be regulated by the length and saturation of phospholipid acyl chains. We therefore evaluated the effect of the acyl chain composition on membrane microviscosity as sensed by DCVJ fluorescence.

We first addressed the variation in saturation using lipid vesicles comprised of DLPC (2× C18:2), DOPC (2× C18:1), and SOPC (C18:0, C18:1). The average membrane viscosity for all compositions is similar within the experimental errors showing that within the analytical error of the method we cannot successfully resolve such subtle differences in viscosity (Fig. 5). The estimated activation energies of the viscosity for DLPC, DOPC, and SOPC (52 ± 13, 54 ± 9, and 68 ± 8 kJ/mol, respectively) show the expected trend DLPC<DOPC<SOPC, although the differences between the activation energies of DLPC and DOPC viscosities are not significant. On the other hand, SOPC shows a relatively high activation energy of 68 ± 8 kJ/mol which is close to the value of the DOPC/cholesterol 6:4 mixture. For chain length variation, we studied membranes composed of phospholipids whose 18- or 16-carbon long acyl chains contained one double bond: SOPC (18:0, 18:1) and POPC (16:0, 18:1). We find that variations in acyl chain structure do not result in statistically significant changes in either the viscosity values (Fig. 6) nor in the viscosity activation energies (53 ± 10 and 68 ± 8 kJ/mol for POPC and SOPC, respectively).

**Figure 5.**
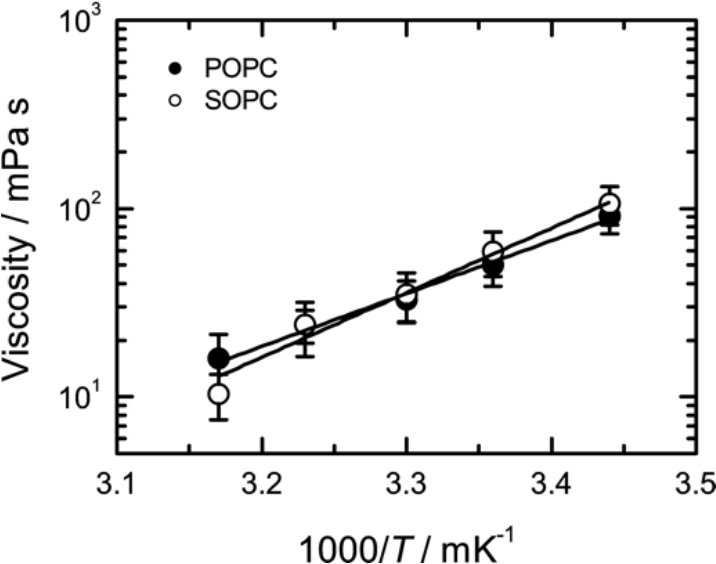
Membrane viscosity variation in response to lipid saturation. Viscosity data are shown along with a fit by the Arrhenius dependence (line). The activation energies of the membrane viscosity are 52 ± 13, 54 ± 9 and 68 ± 8 kJ/mol for DLPC, DOPC and SOPC liposomes, respectively.

**Figure 6.**
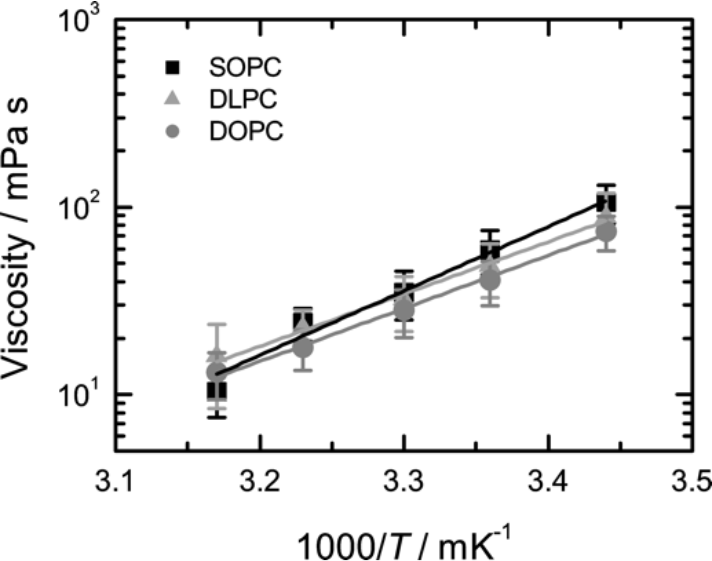
Viscosity of SOPC and POPC lipid membrane (symbols) along with their fits using the Arrhenius law (lines). Activation energies are 53 ± 10 and 68 ± 8k J/mol for POPC and SOPC vesicles, respectively.

Taken together, our results demonstrate that the microviscosities of lipid membranes as reported by DCVJ fluorescence, do not show a pronounced dependence on either the length or saturation of acyl chains.

### 4. Localization of DCVJ in lipid bilayer

The assay presented in this work reports the viscosity of an immediate surrounding of the fluorescent probe. Therefore, the results obtained using this approach should reflect the particular localization of the dye in the membrane. A lot can be learned already from the position of the fluorescence emission peak of DCVJ in the membrane. It has been previously shown that the position of the emission peak of DCVJ is correlated with the solvent polarity.^[15]^ The emission peak of DCVJ in lipid membranes is very close to that of its fluorescence in neat glycerol, suggesting that DCVJ is located in the vicinity of the glycerol backbone of phospholipids rather than around the terminal methyl moiety of acyl chains. Further, high sensitivity of DCVJ to the cholesterol content of the lipid membrane indicates its preferential localization in the proximity of membrane cholesterol, which typically resides next to the glycerol backbone in the direction of the acyl chains.^[28]^ Moreover, this localization of the dye is stable as we did not detect noticeable changes in emission spectra of DCVJ for different membrane compositions and temperatures (data not shown).

Taken together, the experimental evidence suggests that DCVJ is stably localized close to the cholesterol pocket of the lipid membrane.

### 5. Measuring viscosity in biologically relevant membranes

To test the applicability of our approach to biological membranes, we measured the temperature dependence of the viscosity of membranes purified from *Mycoplasma mycoides*, one of the simplest living organisms (Fig. 7).^[29]^ It has been previously shown that at temperatures above the growth conditions the lipid membrane of microorganisms is in Ld state,^[30,31]^ At temperatures lower than the growth temperature, the membranes of microorganisms are known to undergo a phase transition to a state characterized by a considerably higher,^[1, 2, 3, 4, 5, 6, 7]^ including a microorganism closely related to *M. mycoides*.^[32]^ Therefore, one should expect this effect to take place also for membranes of *M. mycoides*. It would be instrumental, therefore, to compare measurements on the *M. mycoides* membranes with two reference lipid mixtures that can model the expected behavior of the bacterial membrane above and below the growth temperature.

**Figure 7.**
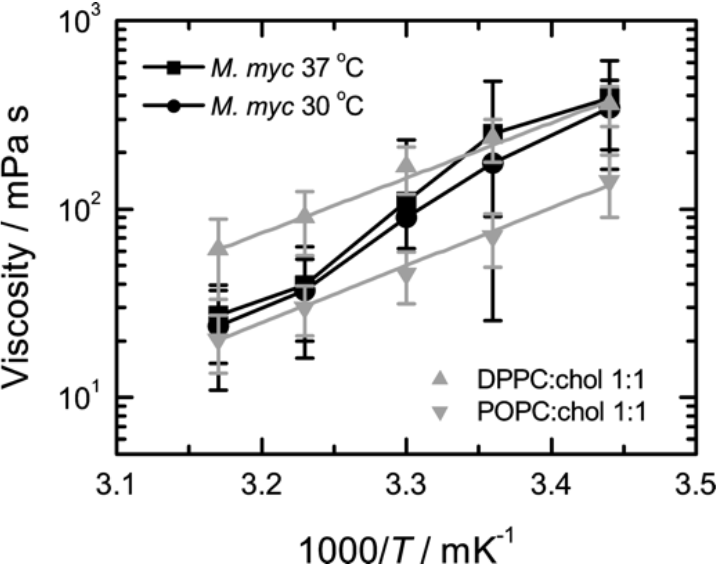
Influence of temperature on viscosity of native lipid membranes from *M. mycoides* grown at 30 and 37 °C (n=5). For comparison, viscosities (symbols) of Ld (POPC/cholesterol 1:1) and Lo (DPPC/cholesterol 1:1) membranes are plotted with corresponding fit of the Arrhenius law (lines).

In order to model the behavior of the membrane at the growth temperature and above, we used the POPC/cholesterol 1:1 lipid mixture, which constitutes vast majority of the native membrane of *Mycoplasma*.^[33]^ This mixture is known to be in the Ld state within the range of temperatures of our study.^[25]^ In contrast, a lipid mixture consisting of DPPC/cholesterol 1:1 that has been shown to be in a more viscous liquid-ordered (Lo) state within the temperature range of our experiments was used as a reference for the more viscous membrane state that the bacterial membrane should be expected below the growth temperature.^[34]^

Comparison of the results for viscosity of *M. mycoides* membranes with those for the artificial lipid mixtures shows that indeed, at temperatures of 37 and 42 °C which are equal or above the growth conditions, bacterial membranes have a very similar viscosity and its temperature dependence to that of the Ld state artificial lipid mixture. At the same time, for the temperatures of 18 and 25 °C which are below the growth temperature, the viscosity of the bacterial membrane approaches the values of the lipid mixture in the Lo phase. Remarkably, the activation energy at the lower end of the temperature range is approximately the same as for the Ld phase at the temperatures. At the same time, a substantial increase in the viscosity activation energy is expected if a transition to the gel phase takes place.^[23]^ This observation suggests that indeed, in agreement with our expectations, the lipid membrane of *M. mycoides* exists in the fluid (liquid-disordered) state at and above the growth temperature, and transforms into a more viscous (liquid-ordered) phase characterized by a higher lipid order at temperatures about 10 degrees below the growth temperature. It also agrees well with the results of Linden et al. showing that cell membranes isolated from *E. coli* exhibit liquid-liquid phase separation upon cooling below the cell growth temperature.^[31]^

According to the concept of homeoviscous adaptation put forward by Sinensky in 1974 the temperature of the phase transition from the fluid phase characteristic of the functional membrane to the more viscous phase-coexistence state should follow the growth temperature of bacteria.^[1]^ This has indeed been previously shown for a large number of microorganisms.^[1, 2, 3, 4, 5, 6, 7]^ Our measurements for membranes of *M. mycoides* grown at two different temperatures, 30 and 37 °C indeed suggest that the expected trend might take place: the transition from the low-viscosity fluid state to the high viscosity phase-separated state is shifted in accordance with the growth temperature. Here, however, we have to point out that the effect is within the experimental error of the method and further experiments are needed to confirm the trend.

## Conclusions

In this work, we showed that using an experimental arrangement based on a standard plate reader capable of simultaneously measuring weak absorbance and relative changes of weak fluorescence signals one can obtain reliable estimates of the lipid membrane viscosity in submicrometer-sized liposomes using a TICT dye – DCVJ. In addition to absolute values of viscosity, we were able to reproducibly measure viscosity activation energies for membranes composed of several different lipid species and their mixtures. We also show that these measurements are compatible with bacterial membranes, and could detect liquid-liquid phase transition in minimal membrane model organism *Mycoplasma mycoides*. Application of a multi-well plate reader would allow one to apply the method to high-throughput bacterial membrane phenotype screening.

## Experimental Section

### Chemicals and Materials

1,2-dioleoyl-*sn*-glycero-3-phosphocholine (dioleoylphosphatidylcholine; DOPC), 1-stearoyl-2-oleoyl-*sn*-glycero-3-phosphocholine (SOPC), 1-palmitoyl-2-oleoyl-glycero-3-phosphocholine (POPC), 1,2-dilinoleoyl-*sn*-glycero-3-phosphocholine (DLPC), and cholesterol were all purchased from Avanti Polar Lipids (Alabaster, AL, USA) and used without further purification. 9-(2,2-dicyanovinyl)julolidine (DCVJ), sodium chloride and 4-(2-hydroxyethyl)piperazine-1-ethanesulfonic acid sodium salt, N-(2-hydroxyethyl)piperazine-N′ -(2-ethanesulfonic acid) sodium salt (HEPES), and anhydrous dimethyl sulfoxide (DMSO) were all obtained from Sigma (St. Louis, MO, USA). Glycerol (spectroscopic grade) and ethylene glycol (spectroscopic grade) were purchase at Alfa Aesar (Ward Hill, MA, USA). For all experiments MiliQ water with resistivity of 18.2 MΩ·cm (25 °C) and TOC below 5 ppb was used. An extruder for liposome preparation was purchased from Avanti Polar Lipids (Alabaster, AL, USA). Whatman® Nuclepore policarbonate filters with pore size of 100 nm were purchased from GE Healthcare (Chicago, IL, USA). Black 96-well plates with transparent bottom (lumox®) were purchased from Sarstedt (Nümbrecht, Germany).

### Liposome preparation

Lipids were pipetted in the form of stock solutions in chloroform (25 mg/mL) into a glass vial, and a thin lipid film was formed on the vial inner surface under the stream of dry nitrogen. Subsequently, the lipid films were kept in vacuum overnight to remove traces of the organic solvent. To form vesicles, the vials containing the lipid films were filled with 10 mM HEPES buffer with 150 mM NaCl at pH 7 and incubated at least 20 °C above the lipid melting temperature for 30 min. Samples were then subjected to 10 cycles of freezing and thawing procedure which was followed by extrusion through a polycarbonate filter with 100 nm pores (10 times). By this means, suspensions of lipid vesicles with the total lipid concentration of 1 mM were formed. Directly before measurements, the vesicle suspensions were diluted to 0.2 mM lipid concentration and DCVJ from a stock solution in DMSO (400 μM) was added to the final concentration of 50 nM resulting in the 400:1 lipid-to-dye ratio. Samples were then incubated for 30 min at 45 °C and 500 rpm using ThermoMixer® (Eppendorf, Wesseling/Berzdorf, Germany). The labelling protocol was tested in a control experiment in which DCVJ was added to lipid solution in chloroform before the formation of the lipid film. The viscosity values we obtained using our method for these samples were identical with those produced using the protocol described above. After incubation, the samples where pipetted into the 96-well plate in the amount of 200 μL per well.

### Preparation of bacterial membranes

*Mycoplasma mycoides* GM12 were grown on SP4 media at 30 and 37 °C and supplemented with Fetal Bovine Serum as a lipid source. Cells were harvested at mid-exponential phase and washed twice in buffer (HEPES 25 mM, NaCl 200 mM, Glucose 1%, pH 7). Washed cells were lysed on an Emulsiflex by passaging three times at 4 bar pressure. The cell lysate was centrifuged at 4000 *g* for 10 minutes to remove non-lysed cells. The lysate was then loaded onto a sucrose step gradient (10%, 30%, 50% w/v) and spun overnight at 4 °C on a Beckman Ti45 rotor at 250000 *g*. A membrane fraction was collected at the 30 %/50 % interface. To remove excess sucrose, the membrane fraction was resuspended in 1.5 mL buffer and pelleted at 70000 *g* for 1 hour. The cell membrane fraction was then resuspended in buffer and stored at −80 °C until analysis.

### Measurement protocol

All spectroscopical measurements were carried out using a SPARK 20M plate reader (Tecan, Grödig, Austria) equipped with a thermostat capable of maintaining the temperature of the sample in the range of 18 - 42 °C with the accuracy of ± 1 °C. The temperature-dependent measurements were carried out at five sample temperatures: 42, 37, 30, 25, and 18 °C starting with 42 °C and subsequently cooling down the sample in steps. Upon reaching a specified temperature, the sample was first incubated for five minutes at 150 rpm using internal sample holder to ensure thermal equilibrium, after which absorption and fluorescence spectra were measured. To reduce evaporation of the samples, the multi-well plate was covered with a lid that was automatically taken off for absorbance measurements and replaced after their completion. At each temperature step, absorption spectra were recorded for each of the wells, after which fluorescence emission spectra from the same wells were collected. Absorbance was measured within the spectral range of 350-550 nm in 2 nm steps with the spectral slit width set to 3.5 nm. Fluorescence emission was measured in the ‘bottom reading mode’ of the setup in the epi-configuration using a 50/50 mirror. Fluorescence excitation and emission wavelengths were selected using the monochromators with the spectral slit widths set to 7.5 nm. Fluorescence was excited at 440 nm using a xenon flash lamp as an excitation source, and the emission spectra were measured in the range of 460-650 nm in 2 nm steps.

### Solution viscosities

Viscosities of glycerol and ethylene glycol at 18, 25, 30, 37, and 42 °C were measured using a Kinexus ultra+ rotational rheometer (Malvern Panalytical, Malvern, Worcestershire, UK) using the cone-plate geometry (1° cone angle, diameter 60 mm) and a temperature stability better than ±0.1 °C. The relative accuracy of the viscosity values was estimated to be better than 3%. Viscosities of glycerol-methanol mixtures were taken from ref. 36.

### Data analysis

Raw data from the plate reader were automatically saved for subsequent processing. For further analysis data were imported into RStudio using home-written script in R language. The ratiometric method described in the present paper requires accurate measurements of the absorption and fluorescence emission of the fluorescent label bound to liposomes. Working at low concentrations typical for the experiments reported here requires that special care should be taken during the analysis of the measured absorbance and fluorescence emission data. Because of the low total concentration of the dye in the volume of the sample (liposome suspension), prior to analysis, the recorded raw spectroscopic data need to be corrected for artefacts in order to extract the pure absorption and fluorescence spectra.

### Absorption spectra

Absorption spectra of DCVJ-labeled liposomes (Fig. 8) consist of the absorbance spectrum of the dye sitting on top of the smooth background sloping down toward longer wavelengths that originates from light scattering by the vesicles in suspension. In our measurement geometry which is based on the use of a multi-well plate, the light beam propagates in the vertical direction and passes through the free liquid-air interface. As a result, the absorption spectrum was found to be additionally affected by the presence of the meniscus. To account for the above-mentioned artefacts, we perform absorbance correction as follows. First, the absorption spectrum of a blank sample containing the same amount of pure buffer was subtracted from the raw absorption spectrum of the sample. This step compensates for the meniscus effect. After this step, the resulting spectrum represents the DCVJ absorption spectrum sitting on top of the smooth background due to light scattering by liposomes. The absorption spectrum of the DCVJ dye represents a well-defined bell-shaped curve, which allowed us to separate the absorption and scattering contributions. In order to do that, the absorption spectrum was recorded within the spectral range wider than the absorption spectrum of DCVJ, and the outermost portions of the measured absorption spectrum (350-374 nm and 526-550 nm) were together fitted by a power law dependence

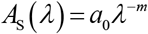

which is known to be a proper phenomenological model to describe the effect of light scattering by liposomes.^[38]^ The scattering background, determined by the fitting routine separately for each well, was then subtracted from the corresponding absorption spectra. By this means, we were able to compensate not only for the above-mentioned phenomena, but also for small absorption offsets caused by the instrument electronics and stray light.

**Figure 8.**
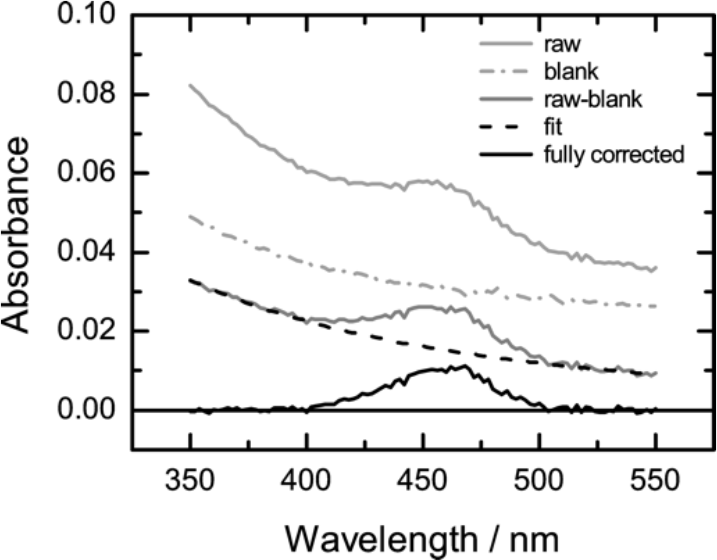
Correction of the absorption spectrum of DCVJ-labeled liposome suspension. Spectra of the sample after consecutive steps of the analysis are shown with solid lines: raw spectrum of DCVJ in liposomes (light gray), result of subtraction of the spectrum of pure buffer from the raw spectrum of the sample (gray) and the resulting “clean” spectrum of DCVJ (black). The dashed line depicts the power-law fit of the scattering background of the sample; the dash-dotted line is the spectrum of the pure buffer. Lipid: DOPC. Temperature: 25 °C.

### Fluorescence spectra

Raw fluorescence spectra of our samples (Fig. 9A) are composed of the DCVJ fluorescence emission spectrum, peak of the Raman scattering of water, and background fluorescence of the 96-well plate. Surprisingly, the background fluorescence of the plate was found to depend on the temperature and showing a progressive increase at its longer wavelength tail upon heating. Furthermore, this effect was systematically stronger at lower rows of the multi-well plate, but reproducible within each of the rows. In order to remove the artefacts from the fluorescence spectra, one well in each row was filled with the pure buffer solution and was used for measuring the blank spectrum. The blank fluorescence emission spectrum (Fig. 9C) was subtracted from the sample spectra (Fig. 9B) to remove the effect of the Raman scattering and background fluorescence of the multi-well plate itself. Knowing that the DCVJ fluorescence emission spectrum in our case goes down to zero at around 640 nm, the constant offset was removed by subtracting the mean signal in the spectral range of 640-650 nm from the spectrum thus resulting in the ‘clean’ fluorescence spectrum of DCVJ in membranes (Fig. 9D).

**Figure 9.**
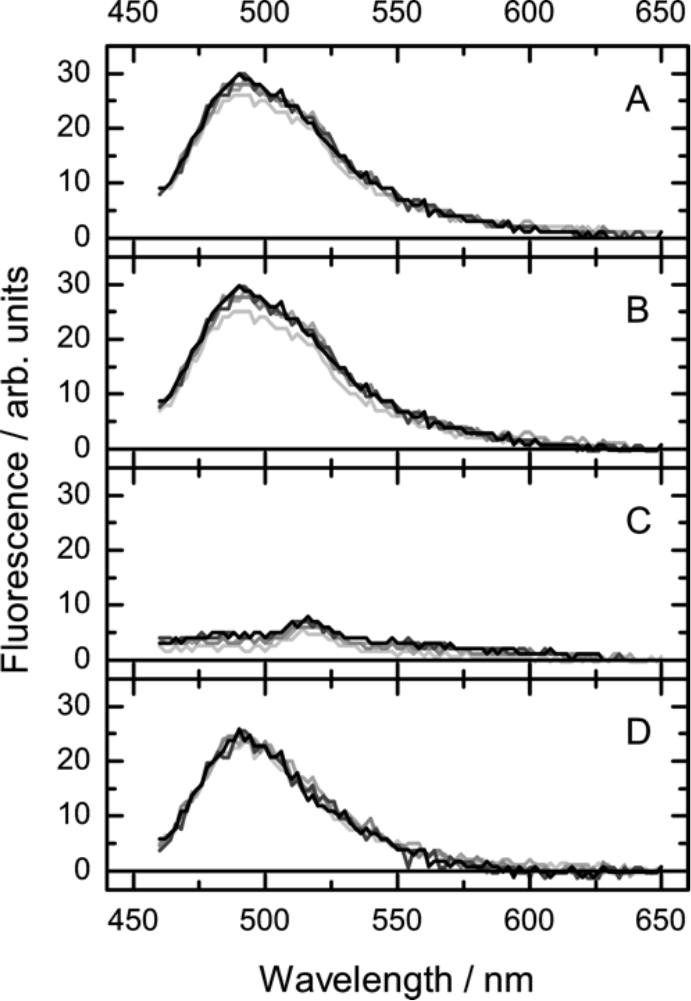
Correction of the fluorescence spectrum of DCVJ-labeled liposome suspension. The figure shows spectra collected from samples located on five different rows of the multi-well plate (each shade of grey corresponds to a different row). (A) raw fluorescence spectra, (B) fluorescence spectra shifted by constant offset, (C) fluorescence background containing the Raman peak of water recorded from the blank sample; (D) ‘pure’ fluorescence spectrum of DCVJ produced by subtracting the spectrum displayed in panel (C) from that in panel (B). Lipid: DOPC. Temperature: 25 °C.

The method is very robust and provides highly reproducible measurement of artefact-free fluorescence spectra under these challenging experimental conditions.

### Estimation of membrane viscosity and its Arrhenius activation energy

The method we describe here is based on the dependence of the fluorescence quantum yield of the DCVJ dye on the microviscosity of its immediate environment.^[16]^ As the experimental determination of the absolute fluorescence quantum yield is very challenging, especially under our experimental conditions, we resort instead to the use of a proxy quantity defined as a ratio of the fluorescence emission intensity and absorbance of membrane-bound DCVJ.

In order to obtain lipid viscosity estimates, we first calculated the ratio *R* = *F*/*A* of the quantities representing the fluorescence intensity (*F*) and absorbance (*A*) of membrane-bound DCVJ.

The quantity *A* was evaluated by integrating the artefact-free absorption spectrum over an interval of wavelengths that was centered on the fluorescence excitation wavelength and had a width corresponding to the spectral slit width in the fluorescence measurements. The quantity *F* was evaluated by integrating the artefact-free fluorescence emission spectrum over the whole spectral range of fluorescence measurements.

Results from three independent replicates (5 analytical replicates each) were grouped together, and the mean values of fluorescence intensity ⟨*F*⟩ and absorbance ⟨*A*⟩, and their standard deviations *σ*_*F*_ and *σ*_*A*_ were calculated. Subsequently, fluorescence-to-absorbance ratio ⟨*R*⟩ = ⟨*F*⟩/⟨*A*⟩ was calculated, and the corresponding standard error was estimated using the error propagation as follows:

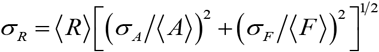

Similar to what has been previously reported,^[15,16]^ the dependence of the fluorescence-to-absorbance ratio *R* of DCVJ on the viscosity of the medium was found to follow the power law (see Figure 2)

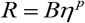

The value of the exponent *p* = 0.53 ± 0.01 was determined by fitting the power law to the dependence of *R* on *η* using a weighted nonlinear least squares routine and was found to be in excellent agreement with the previous findings.^[15]^ The value of the constant prefactor *B* depends on the particular instrument and measurement settings (which were kept constant in our measurements), and is therefore not reported here. Reverting this relationship allows one to obtain an estimate of the microviscosity of the DCVJ environment based on the measured value of the fluorescence-to-absorption ratio *R*. The uncertainties in the estimates of the microviscosity were calculated using the error propagation as follows:

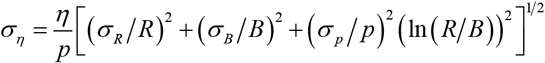

To obtain the Arrhenius activation energies of the viscosity, the temperature dependences of the estimated viscosities of the samples were analysed using the Arrhenius law

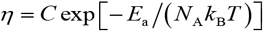

where *E*_a_ is the activation energy, *T* is the absolute temperature, *N*_A_ is the Avogadro number, *k*_B_ is the Bolzman constant, and *C* is the numerical prefactor (irrelevant for our study). The analysis was based on a weighted nonlinear least squares routine that took into account the uncertainties in the viscosity values and provided the best estimate of the fitting parameters along with the estimates of their uncertainties.

### Partitioning of DCVJ in lipid membrane

The present method is based on the assumption that only the dye molecules which partition into the lipid membrane contribute to the measured ratio *R*. Since DCVJ does not fluoresce in the aqueous environment but still is capable of absorbing light, the presence of non-membrane-bound dye molecules in the sample would lead to underestimation of *R*. Based on the chemical structure of the dye, its strong affinity toward the non-polar lipid environment should be expected, but still requires an experimental verification. Conveniently, our setup allows us to monitor both the dye and lipid vesicle content in the sample via absorption measurements (see above). Namely, the smooth background due to scattering of light by liposomes is proportional to their concentration, whereas the DCVJ absorption peak ‘sitting’ on top of that background is proportional to the dye content in the sample. If the dye predominantly partitions in the lipid membrane, then removal of liposomes from the sample should result in the proportional decrease in the liposome scattering contribution and the DCVJ absorption peak. To test whether this is indeed the case, we carried out a series of consecutive ultracentrifugation steps. At each step, the sample was subjected to ultracentrifugation at 186000 *g* for 45 min, after which the supernatant depleted in the content of lipid vesicles was carefully collected, and was used partly for absorbance measurements and partly for the next ultracentrifugation step. We carried out three consecutive ultracentrifugation steps, and found that after each step both the liposome scattering contribution and DCVJ absorbance dropped in a correlated manner. As an additional step, we subjected our samples to ultrafiltration (Amicon Ultra 10K, Merck KGaA, Darmstadt, Germany) which was expected to remove the liposomes from the sample, but not affect the presence of the small dye molecules potentially dissolved in the buffer solution. The samples subjected to ultrafiltration showed absorption spectra indistinguishable from the base line, indicating the absence of both the liposomes and DCVJ in the sample. A linear fit of the data set comprised of the measurements on a sample prepared in a standard manner, three samples produced from the original sample by consecutive ultracentrifugation steps, and a sample produced from the original sample by ultrafiltration, shows an intercept statistically indistinguishable from zero (Fig. 10). These results fully justify the assumption that the presence of free, non-membrane-bound, DCVJ dye in our samples can be neglected.

**Figure 10.**
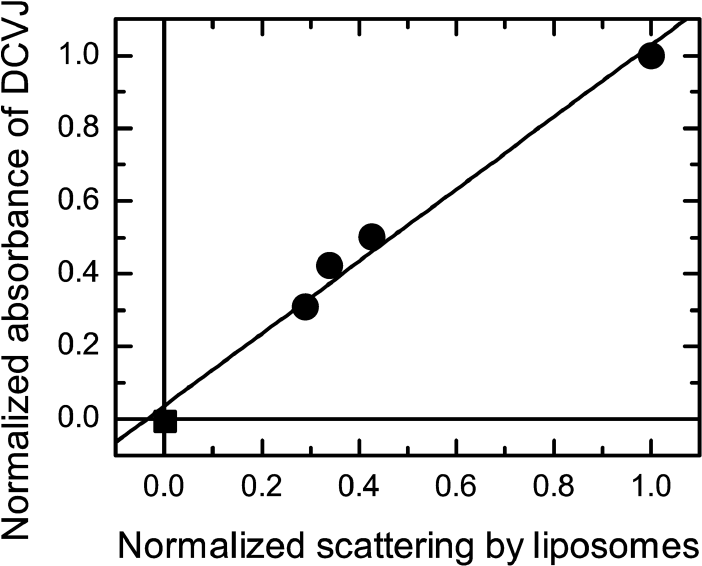
Normalized absorbance of the DCVJ as a function of the normalized contribution of scattering of light on liposomes to the absorption spectrum upon progressive removal of liposomes from the original sample. Circles: original sample and samples produced from it by three consecutive ultracentrifugation steps. Square: sample produced by ultrafiltration of the original sample. Lipid: DOPC. Temperature: 25 °C.

## Acknowledgements

Authors are grateful to Marén Gnädig for her technical support and membrane isolations. This work was supported by the B CUBE of the Technische Universität Dresden, a German Federal Ministry of Education and Research (BMBF) grant (to J.P.S., project # 03Z22EN12), and a VW Foundation ‘Life’ grant (to J.P.S., project # 93090). E.P.P. acknowledges the financial support by the DFG within the SFB 1032 TP B01 and A09.

---

We developed a robust and financially accessible quantitative method to measure microviscosity of lipid membranes, which is suitable for highthroughput screening. The approach is based on the simultaneous measurements of absorbance and fluorescence emission of a fluorescent dye and allows for monitoring of the temperature dependence of viscosity in bacterial membranes.

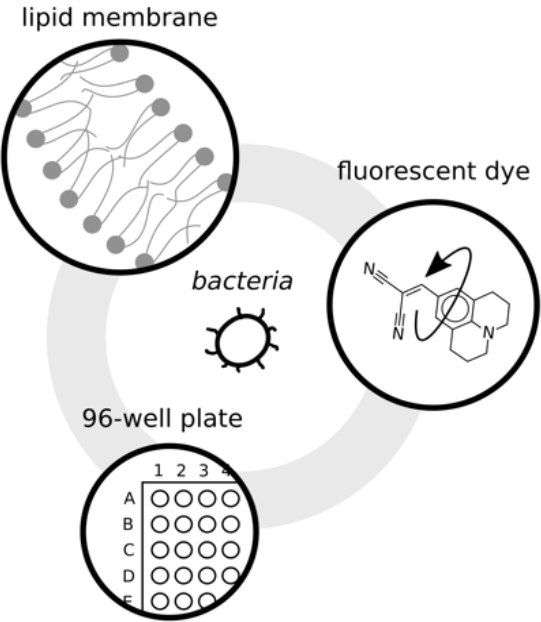

